# Explainable Deep Generative Models, Ancestral Fragments, and Murky Regions of the Protein Structure Universe

**DOI:** 10.1101/2022.11.16.516787

**Authors:** Eli J. Draizen, Cameron Mura, Philip E. Bourne

## Abstract

Modern proteins did not arise abruptly, as singular events, but rather over the course of at least 3.5 billion years of evolution. Can machine learning teach us how this occurred? The molecular evolutionary processes that yielded the intricate three-dimensional (3D) structures of proteins involve duplication, recombination and mutation of genetic elements, corresponding to short peptide fragments. Identifying and elucidating these ancestral fragments is crucial to deciphering the interrelationships amongst proteins, as well as how evolution acts upon protein sequences, structures & functions. Traditionally, structural fragments have been found using sequence-based and 3D structural alignment approaches, but that becomes challenging when proteins have undergone extensive permutations—allowing two proteins to share a common architecture, though their topologies may drastically differ (a phenomenon termed the *Urfold*). We have designed a new framework to identify compact, potentially-discontinuous peptide fragments by combining (i) deep generative models of protein superfamilies with (ii) layerwise relevance propagation (LRP) to identify atoms of great relevance in creating an embedding during an all_*superfamilies*_ × all_*domains*_ analysis. Our approach recapitulates known relationships amongst the evolutionarily ancient small *β*-barrels (e.g. SH3 and OB folds) and amongst P-loop–containing proteins (e.g. Rossmann and P-loop NTPases), previously established via manual analysis. Because of the generality of our deep model’s approach, we anticipate that it can enable the discovery of new ancestral peptides. In a sense, our framework uses LRP as an ‘explainable AI’ approach, in conjunction with a recent deep generative model of protein structure (termed *DeepUrfold*), in order to leverage decades worth of structural biology knowledge to decipher the underlying molecular bases for protein structural relationships—including those which are exceedingly remote, yet discoverable via deep learning.

## 1 Introduction

Historically, protein structural evolution has been studied via painstaking visual inspection and manual analyses of structures, including a heavy reliance on comparisons built upon 3D superposition/alignment of atomic coordinates. Indeed, visualizing protein structures and their 3D alignments using graphical tools such as ‘ribbon diagrams’ enabled many landmark discoveries in biology and medicine, largely because these diagrams simplify the inordinately complicated geometric structures of proteins into something that is comprehensible by the human brain [1]. However, simplified cartoon representations ignore the high-dimensional, largely biophysical/physicochemical feature space of all the atoms defining a protein, and an over-reliance on simplified, static representations can limit us to seeing *a* structure of a particular protein as being *the* structure—i.e., we fall prey to viewing as important only one particular geometric representation or structural conformation, versus the true statistical ensemble of thermally-accesible states that an actual protein structure samples in reality. In short, conventional schemes for conceptualizing and analyzing protein structure relationships are not without limitations, and can cause us to miss phylogenetically remote (deeply ancestral) relationships. Some have referred to this pitfall as the ‘curse of the ribbon’ [1]. Modern deep learning–based methods enable fundamentally new representations of proteins and their sequence/structure/function relationships—for example, as lower-dimensional embeddings that incorporate biophysical properties (e.g., electrostatics) alongside 3D-coordinate data (i.e., geometry), sequence information and residue profiles, and so on. All of this, in turn, might finally lift the curse.

While many new sequence-based deep learning methods, based on large language models [2, 3, 4, 5], can identify more remote similarities than can hidden Markov models (HMMs) or classic sequence-comparison algorithms (e.g., BLAST), larger structural rearrangements and permutations, such as occur on evolutionary timescales, are still difficult to detect. If one views the protein universe through the lens of a hierarchical classification scheme such as CATH [6], most new homologous sequences identified by these methods would be located within the same homologous superfamily or topology strata—i.e., there are bounds on how remote a homology can be detected by existing methods. Thus, sets of potentially distantly-related proteins, with similar architectures yet different topologies, are generally missed [7]. Indeed, it was recently proposed that there might exist a new, bona fide level of structural granularity, lying between the architecture and topology levels (the latter of which is synonymous with a given protein’s *fold*); termed the ‘Urfold’, this provisional new level would naturally allow for 3D fragments that are spatially compact yet potentially discontinuous in sequence [7], such as may be the case with ancient, deeply ancestral fragments. An example of an urfold can be seen in Fig 1, highlighting two widespread classes of phosphate-binding loop (PBL)– containing proteins, namely the Rossmann fold-containing proteins and P-loop NTPases.

**Figure 1:**
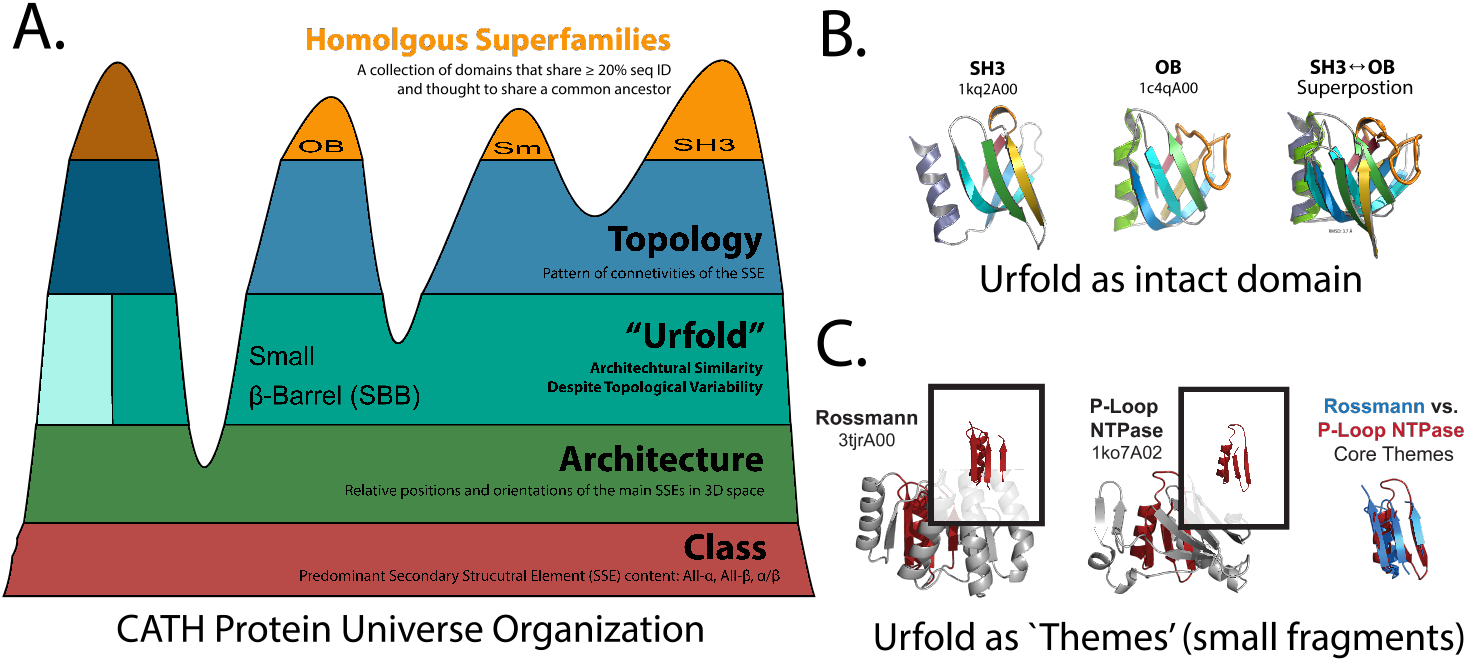
Urfolds Represent Architectural Similarity Despite Topological Variability. A) CATH hierarchically organizes the protein universe into strata including Class, Architecture, Topology, and Homologous Superfamily. We hypothesize that an ‘Urfold’ strata between Architecture and Topology. B) SH3 and OB share a small *β*-barrel urfold. C) The Rossmann and P-loop NTPases both contain a common core theme of three *β*-strands connected to an *α*-helix by a glycine-rich loop; the loop is of functional importance by virtue of binding phosphorylated ribonucleoside ligands, be they substrates, cofactors, or otherwise. However, the Rossmann fold is permuted such that one *β*-strand is no longer nearby in sequence and another *β*-strand has reversed direction. These gross structural differences have led biologists to incorrectly classify these ancient folds as being unrelated. We propose bridging these two folds via a new ‘PBL’ urfold [12, 7].

The Urfold model of protein structure—and, thus, protein structural interrelationships—arose by noticing the striking structural and functional similarities among deeply-divergent collections of domains from the SH3 and OB superfamilies [8]. Those superfamilies all contain a small *β*-barrel (SBB) domain, composed of five *β*-strands, but the topologies/connections between the strands have been permuted such that these proteins often share less than 20% sequence identity between one another (i.e., below the classic ‘twilight zone’ of inferring homology as above-random similarity between two sequences). Despite having permuted strands, the architecturally-identical SBBs are often involved in nucleic acid metabolic pathways, and many of them oligomerize via residue interactions amongst similar edge-strands [8]. Recently, others have proposed that the SH3 and OB share a common ancestor that diverged via a process called ‘Creative Destruction’ [9, 10]. Notably, the SH3 and OB are two of the most ancient and widespread protein folds, and they permeate most information-storage and information-processing pathways in cellular life, from DNA replication to transcription of DNA→RNA and translation of RNA→protein [9, 11].

Another prominent example of an urfold can be seen in the phosphate-binding loop (PBL) containing proteins, which include the Rossmann and P-loop NTPases superfamilies [12]. Both of these superfamilies contain a small protein fragment of three *β*-strands that contact a single *α*-helix, with these structural elements linked by a phosphate-binding glycine-rich loop (Fig. 1). Most domains from the PBL urfold are quite large, i.e. they feature many additional secondary structural elements (SSEs) beyond the core, and they do not oligomerize; rather, they have a large central cavity for ligand-binding. While Rossmann and PBL proteins are known to bind a diverse repertoire of ligands and catalyze many different types of reactions, they predominately bind phosphorylated nucleotides and similar compounds, with the ligand’s phosphate groups primarily interacting with the protein’s phosphate-binding loop [13].

In addition to similarity at the full-domain level, the Urfold model allows for sub-domain–level structural fragments that may be discontinuous in sequence. That is, the Urfold extends a CATH-like hierarchical representational scheme of the protein universe by allowing for (conserved) spatial constellations of short peptides, perhaps like the *Ancestral Peptides* [14] or *Themes* [15] that have been thought to underlie the structural evolution into larger domains. Note that algorithms which ?exibly allow for discontinuous fragments are making a resurgence, e.g. as in the *Geometricus* approach [16] for constructing and analyzing structural embeddings.

In a recent study that developed a deep generative approach to protein structural relationships, using the Urfold model of protein structure in a framework called *DeepUrfold*, 20 superfamily-specific, sparse 3D-CNN variational autoencoders (VAEs) were trained for 20 different, highly-populated CATH superfamilies [17]. These DeepUrfold-trained models were shown to be agnostic to topology, as architecturally-similar SH3/OB proteins with artificially-constructed loop permutations yielded similar evidence lower bound–based (ELBO) scores; most significantly, applying community-detection methods (as stochastic block models) to the patterns of ELBO similarities led to the SH3 and OB domains clustering into similar groupings (with some intermixing). All of those findings were consistent with the prediction that the SH3 and OB comprise a distinct urfold (in this case, the SBB).

This paper explores—and seeks to begin *explaining*—the models from [17] in more depth, by applying an approach known as layer-wise relevance propagation (LRP). In principle, explainable AI techniques such as LRP can be used to understand which atoms in the input structure are ‘important’, based on their spatial locations and biophysical properties (and, really, any other sorts of features that one encodes in the model). In our all_*super families*_ × all_*domains*_ analysis, we consider the ELBO likelihood of a domain *x* under under *DeepUrfold* VAE models M_*i*_ and M_*j*_, for superfamilies *i* and *j* respectively. Functionally conserved regions shared between M_*i*_ and M_*j*_ would be expected to positively affect the likelihood under both models and therefore should have relatively high LRP scores. Focusing on the two specific urfolds described above—i.e., the SBB domain and the PBL-containing proteins—here we show how LRP can be used to identify cross-model functionally important atoms; achieving that task, in turn, offers a foundation for identifying and characterizing new discontinuous fragments or ancestral peptides.

## 2 Results

### 2.1 The small *β*-barrel (SBB) domain

We first investigated the SH3-specific (CATH 2.30.30.100), DeepUrfold-derived VAE model. This model was trained using all energy-minimized domain structures from the SH3 superfamily, along with hand-crafted biophysical features, as described in [17]. We first attempted to subject representative SH3 domains through the SH3 model and calculated relevance scores during backpropagation. Promisingly, all of the residues that were previously identified (manually) as being important in the SBB’s “conserved hydrophobic core” [8] were labelled as relevant according to our LRP calculation (Fig. 2B). Next, we found that a specific, highly-conserved glycine in the second *β*-strand, known from years of manual analysis as being conformationally important in allowing the strand to severely bend [18], was deemed to be ‘relevant’ (Fig. 2C). Finally, we show that many contacts between strands *β*4 and *β*5, such as comprise the subunit interface in the ‘Sm’ variety of SH3-based oligomeric rings, are also labelled as important (Fig. 2D). From these initially promising results, we suspect that LRP can identify functionally important atoms, such as are learned as part of the latent space in the DeepUrfold-based superfamily models.

**Figure 2:**
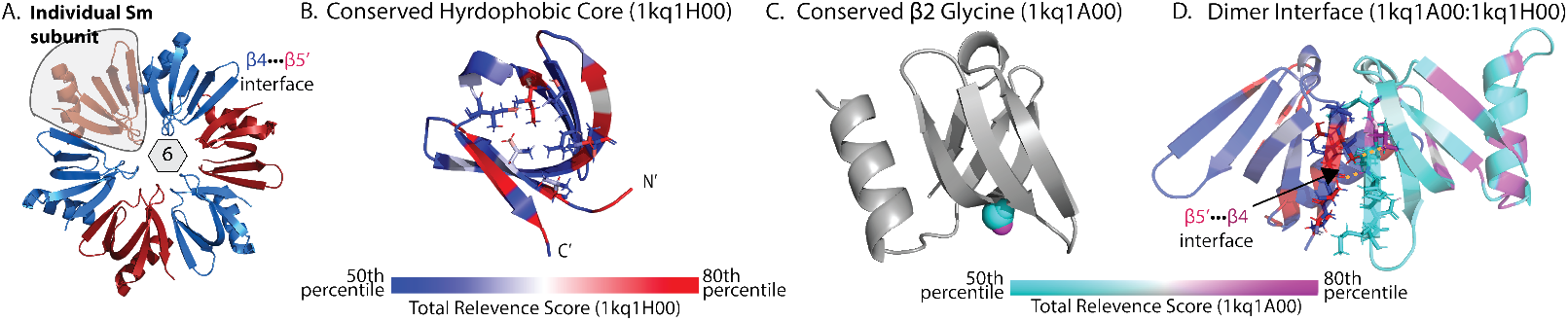
LRP Identifies Conserved and Structurally Important Regions in SH3 Domains. A) SH3 domains, and specifically those of the Sm/Sm-like proteins, tend to self-assemble into oligomeric rings of *n* = 5, 6, or 7 SH3 subunits [18]. B) All SH3 domains, as well as other domains of the SBB urfold, have a conserved hydrophobic core [8]. LRP identifies residues in the core as being 80th percentile or greater, displayed here using a spectrum from blue→white→red, for the range of 50th to 80th percentile, and for all of the 1kq1H00 relevance scores. C) The phylogenetically conserved, structurally critical *β*2 glycine [18] is detected by LRP. The color scale is shown for 1kq1A00 (with a cyan→ white → magenta spectrum), mapped across the range 50th to 80th percentile for all of the 1kq1A00 relevance scores. D) By manual/visual inspection of the 1kq1H00:1kq1A00 dimer interface, we can see that important atoms from both 1kq1H00 roughly align with important atoms in 1kq1A00 (yellow dashed lines). Note that 1kq1H00 is shown on the same color scale as in (B), and 1kq1A00 is shown with the same color scale as in (C).

### 2.2 Phosphate-binding Loop (PBL)–Containing Proteins

We next tested the outcome of an all_*superfamily*_ × all_*domain*_ approach for the PBL urfold. That is, we trained two separate DeepUrfold VAE models, one for the canonical Rossmann Fold (3.40.50.720) and another for P-loop NTPases (3.40.50.300), and then we subjected representative domains from both superfamilies through both VAEs. Given the biochemical importance of phosphate binding, we examined residues that are known to specifically bind phosphates: namely, the glycine-rich loop and the Walker B motif’s aspartic acid on one of the edge strands of the PBL theme [12]. For all four combinations of domain× model—i.e., (i) `Rossmann domain→ Rosmann model`, (ii) `Rossmann domain → P-Loop NTPase model`, (iii) `P-Loop NTPase domain → Rossmann model`, and (iv) `P-Loop NTPase domain→ P-Loop NTPase model`—we find that LRP correctly identifies the glycine-rich loop and Walker B Asp with relevance scores >=75th percentile, shown in Fig 3. Because the important atoms are correctly predicted regardless of the DeepUrfold model, even for a model that is trained on domains annotated from a different CATH superfamily, we expect that these important residues, shared by both the Rossmann and PBL families, can be used to identify common fragments that comprise a joint Rossmann/PBL urfold. LRP-?agged atoms can be viewed, in a very real sense, as identifying the ‘defining’ regions for a particular urfold.

**Figure 3:**
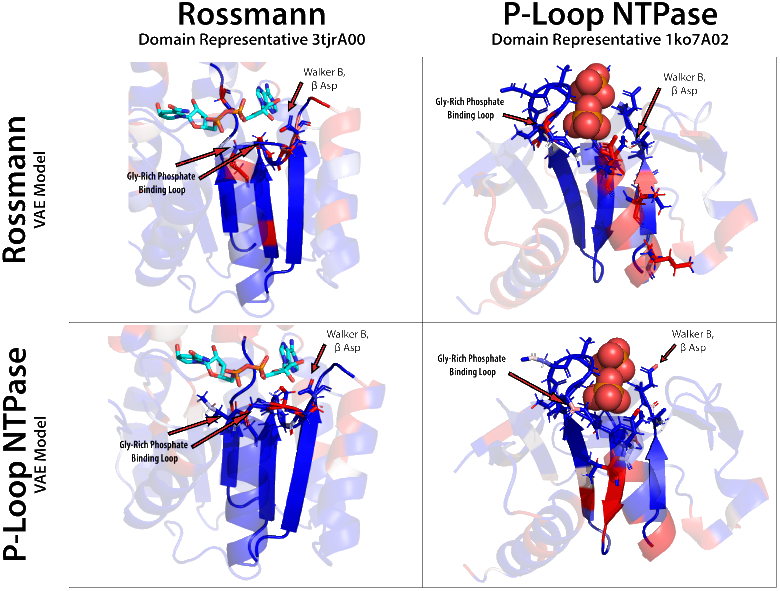
Important Atoms in the Phosphate Binding Loop Urfold Identified via LRP. We subjected representative domains from the Rossmann (3tjrA00) and P-Loop NTPase (1ko7A02) superfamilies through each VAE model trained on all members of Rossmann *and* P-Loop NTPases respectively. Relevance scores are displayed on a spectrum from blue → white → red, using a range of 50th to 80th percentile of a given structure. Key atoms from residues that have been previously shown to be important in bridging these folds, namely the Walker B Asp motif and the glycine-rich loop [12], are selected by LRP having an relevance score >=75th percentile. Included ligands highlight phosphate-binding regions.

## 3 Methods

### 3.1 *DeepUrfold-Explain* and VAE Model

In a recent paper that introduced a *DeepUrfold* framework, the authors developed: (i) a preprocessed dataset, based on CATH superfamilies, that includes biophysical properties for each atom along with energy-minimized domain structures; and (ii) superfamily-specific sparse 3D-CNN VAEs [17]. Energy-minimized domain structures from a single superfamily were voxelized using a *k*D-tree to map atoms into 1Å^3^ voxels in an overall 264^3^Å^3^ discretized volume; 3D structural models were rotated randomly by sampling the *SO*(3) group to train a VAE model, modified from [19, 20], yielding superfamily-specific models. Each voxel is annotated with the biophysical properties of atoms that intersect it (see the features listed in Fig. 4). Each VAE was trained using CATH’s 30% sequence identity clusters, as defined by CATH for each superfamily, to create train (≈ 80%), test (≈ 10%), and validation (≈10%) splits. Hyperparameters used to construct and refine the VAE were tuned using Weights & Biases [17].

**Figure 4:**
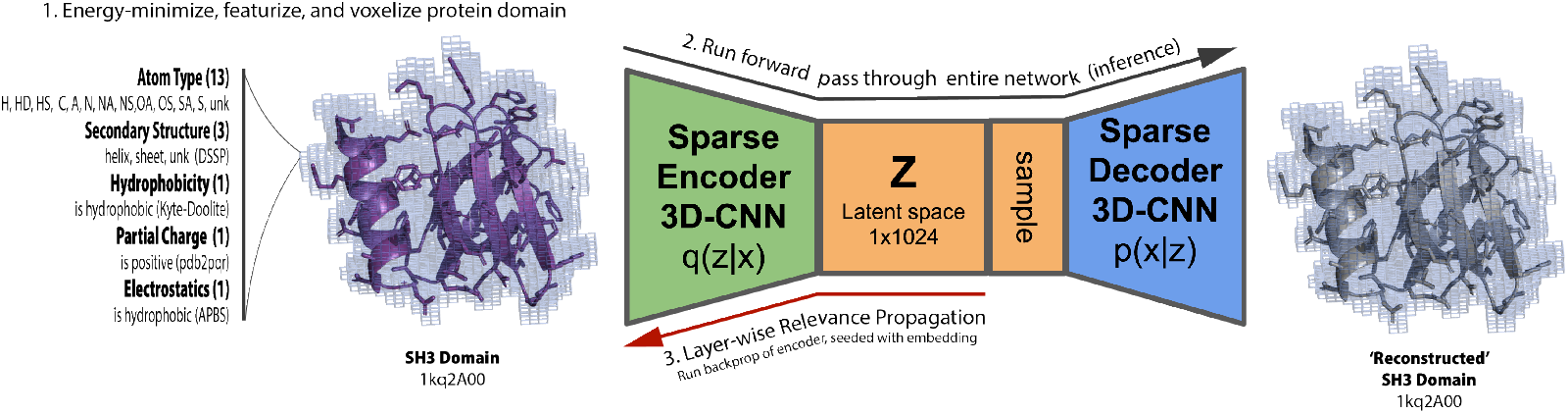
*DeepUrfold-Explain* Identifies Important Atoms in Input Structures, via LRP of Superfamily-specific VAEs. Relevant atoms are predicted for a given protein domain by: 1) obtaining an energy-minimized, featurized domain structure from a pre-calculated dataset and voxelized; 2) running the inference stage of pre-trained superfamily-specific VAE for the given domain; and finally, 3) running LRP during a backwards pass of only the encoder module, starting with the embedding of a given domain.

### 3.2 Layer-wise Relevance Propagation (LRP)

The LRP algorithm defines a Relevance Score as 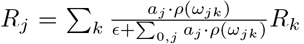, where *j* denotes a neuron in the current layer, *a*_*j*_ are the activations from the current layer, *ρ* is the LRP propagation ‘rule’, and *ω*_*jk*_ are the weights from the previous layer. LRP starts with the embedding of a given domain in a backwards pass. For layer *j*, a forward pass is run with the same data that was used as input to the layer *j* (the denominator) which is compared to the Relevance value from the previous layer, *R*_*k*_. A backwards pass is then run using the data from the relevance-weighted forward pass. Finally, the value from the backwards pass is compared to the output of the original input of the current layer [21].

We follow [22] and use rules LRP-0 for the lowest 60% of layers, LRP-*ϵ* (*ϵ*=0.25) for the middle 20%-60% of layers, and LRP-*γ* (*γ*=0.25) for the top 20% of layers. LRP-0 is the base rule, where *ρ*(*x*) = *x* and *ϵ* = 0, which finds the contributions of each neuron to the final activation. LRP-*ϵ* is the base rule where *ρ*(*x*) = *x*, but *ϵ >* 0, which helps remove noise and sparsify the explanations in terms of the input. LRP-*γ* sets *ρ*(*x*) = *x* + *γx*^+^, where *x*^+^ only includes positive relevance scores (all others set to 0) and *ϵ* = 0, which is used to remove negative contributions.

Finally, we create a total relevance score by aggregating all relevance scores in a voxel; this was done by summing the relevance scores for every feature in that voxel. We then map voxels*↔*atoms by taking the average total relevance score from all of the voxels that intersect a given atom based on a *k*D-tree, the radius of which is set to the size of the particular atoms (in terms of van der Waals radius).

We adapted PyTorchLRP from [23] to add MinkowskiEngine layers [19] and regularization [24].

### 3.3 Cross-Model Fragment Identification

We subjected 2,674 representative domains from 20 different CATH superfamilies to 20 superfamily-specific VAEs, saving all LRP results. For a given structure, residues containing any atom lying in the ≥ 75^th^ percentile were extracted to create a set of 53,480 (dis-)continuous fragments. For each community identified with Stochastic Block Modelling of the bipartite graph formed from the all_*superfamilies*_× all_*domains*_ approach [17], we used the foldseek code [25] to cluster all LRP structures from domains present in each community that were processed through all superfamilies represented by the community (using TM-Align for global alignment). We select the LRP structure cluster representative from the most populated cluster in each community, resulting in a final list of top-20 “potential urfolds”.

## 4 Conclusion

Machine learning for proteins is extremely difficult, partly due to the fact that all proteins are related via evolution [26]. As one can imagine, it is critical to know if a given model accurately represents reality or is giving spurious results. Explainable AI techniques, such as LRP, help alleviate these problems by allowing us to compare known biophysical properties of a given protein to a model’s prediction—that is, we can bring the expertise from a scientific domain to bear in assessing the performance of the ML method. Here, we have shown that LRP correctly identifies structurally important and conserved atoms in SH3 domains, implying that the model is learning superfamily-specific features. Because the models are intentionally topology-agnostic, we were also able to show that LRP can find important atoms from structures that exhibit the Urfold principle of “*architectural similarity despite topological variability*”—specifically, in this work the phosphate binding loops in Rossmann-fold proteins and P-Loop NTPases. In the future, we plan to identify and verify more common fragments and ancestral peptides by aligning and clustering ‘important’ regions from the cross-model fragments identified by LRP, while comparing them to known databases of potentially discontinuous fragment libraries (e.g., from shapemers [27], Fuzzle2.0 [28], ancestral peptides [14], themes [15], or TERMs [29]).

## 5 Appendix

### 5.1 Variational Autoencoders (VAEs)

Each VAE model learns a normal distribution for each superfamily using the ‘reparameterization trick’ to allow for backpropagation through random variables. This makes only the mean (*µ*) and variance (*σ*) differentiable, while sampling from the normally distributed random variable (*𝒩* (0, **I**)). That is, the latent variable posterior **z** is given by **z** = *µ* + *σ*⊙ *𝒩* (0, **I**), where ⊙ denotes the Hadamard (element-wise) matrix product and *𝒩* is the ‘auxiliary noise’ term [30]. See [17] for a discussion of this topic in the context of DeepUrfold.

## References

[1] Philip E. Bourne, Eli J. Draizen, and Cameron Mura. The curse of the ribbon. PLoS Biology, Accepted 2022.

[2] Vamsi Nallapareddy, Nicola Bordin, Ian Sillitoe, Michael Heinzinger, Maria Littmann, Vaishali Waman, Neeladri Sen, Burkhard Rost, and Christine Orengo. CATHe: Detection of remote homologues for CATH superfamilies using embeddings from protein language models. bioRxiv, 2022.

[3] Michael Heinzinger, Maria Littmann, Ian Sillitoe, Nicola Bordin, Christine Orengo, and Burkhard Rost. Contrastive learning on protein embeddings enlightens midnight zone. NAR Genomics and Bioinformatics, 4(2), 06 2022. lqac043.

[4] Tymor Hamamsy, James T. Morton, Daniel Berenberg, Nicholas Carriero, Vladimir Gligorijevic, Robert Blackwell, Charlie E. M. Strauss, Julia Koehler Leman, Kyunghyun Cho, and Richard Bonneau. TM-Vec: Template modeling vectors for fast homology detection and alignment. bioRxiv, 2022.

[5] Tristan Bepler and Bonnie Berger. Learning the protein language: Evolution, structure, and function. Cell Systems, 12(6):654–669.e3, June 2021.

[6] Ian Sillitoe, Nicola Bordin, Natalie Dawson, Vaishali P Waman, Paul Ashford, Harry M Scholes, Camilla S M Pang, Laurel Woodridge, Clemens Rauer, Neeladri Sen, Mahnaz Abbasian, Sean Le Cornu, Su Datt Lam, Karel Berka, Ivana Hutařová Varekova, Radka Svobodova, Jon Lees, and Christine A Orengo. CATH: increased structural coverage of functional space. Nucleic Acids Research, 49(D1):D266–D273, November 2020.

[7] Cameron Mura, Stella Veretnik, and Philip E. Bourne. The Urfold: Structural similarity just above the superfold level? Protein Science, 28(12):2119–2126, November 2019.

[8] Philippe Youkharibache, Stella Veretnik, Qingliang Li, Kimberly A. Stanek, Cameron Mura, and Philip E. Bourne. The small β-barrel domain: A survey-based structural analysis. Structure, 27(1):6–26, January 2019.

[9] Claudia Alvarez-Carreño, Petar I Penev, Anton S Petrov, and Loren Dean Williams. Fold evolution before LUCA: Common ancestry of SH3 domains and OB domains. Molecular Biology and Evolution, 38(11):5134–5143, August 2021.

[10] Claudia Alvarez-Carreño, Rohan J Gupta, Anton S. Petrov, and Loren Dean Williams. The evolution of protein folds by creative destruction. bioRxiv, 2022.

[11] Vishal Agrawal and Radha KV Kishan. Functional evolution of two subtly different (similar) folds. BMC Structural Biology, 1(1):1–6, 2001.

[12] Liam M Longo, Jagoda Jabłońska, Pratik Vyas, Manil Kanade, Rachel Kolodny, Nir Ben-Tal, and Dan S Tawfik. On the emergence of P-Loop NTPase and Rossmann enzymes from a beta-alpha-beta ancestral fragment. eLife, 9, December 2020.

[13] Kirill E. Medvedev, Lisa N. Kinch, R. Dustin Schaeffer, and Nick V. Grishin. Functional analysis of Rossmann-like domains reveals convergent evolution of topology and reaction pathways. PLOS Computational Biology, 15(12):e1007569, December 2019.

[14] Vikram Alva, Johannes Söding, and Andrei N Lupas. A vocabulary of ancient peptides at the origin of folded proteins. eLife, 4, December 2015.

[15] Sergey Nepomnyachiy, Nir Ben-Tal, and Rachel Kolodny. Complex evolutionary footprints revealed in an analysis of reused protein segments of diverse lengths. Proceedings of the National Academy of Sciences, 114(44):11703–11708, October 2017.

[16] Janani Durairaj, Mehmet Akdel, Dick de Ridder, and Aalt D J van Dijk. Geometricus represents protein structures as shape-mers derived from moment invariants. Bioinformatics, 36(Supplement_2):i718–i725, December 2020.

[17] Eli J. Draizen, Stella Veretnik, Cameron Mura, and Philip E. Bourne. Deep generative models of protein structure uncover distant relationships across a continuous fold space, 2022.

[18] Cameron Mura, Peter S. Randolph, Jennifer Patterson, and Aaron E. Cozen. Archaeal and eukaryotic homologs of Hfq: A structural and evolutionary perspective on Sm function. RNA Biology, 10(4):636–651, April 2013.

[19] Christopher Choy, JunYoung Gwak, and Silvio Savarese. 4D spatio-temporal convnets: Minkowski convolutional neural networks. pages 3075–3084, 2019.

[20] JunYoung Gwak, Christopher B Choy, and Silvio Savarese. Generative sparse detection networks for 3D single-shot object detection. 2020.

[21] Alexander Binder, Grégoire Montavon, Sebastian Bach, Klaus-Robert Müller, and Wojciech Samek. Layer-wise relevance propagation for neural networks with local renormalization layers. 2016.

[22] Grégoire Montavon, Alexander Binder, Sebastian Lapuschkin, Wojciech Samek, and Klaus-Robert Müller. Layer-wise relevance propagation: An overview. pages 193–209, 2019.

[23] Moritz Böhle, Fabian Eitel, Martin Weygandt, and Kerstin Ritter. Layer-wise relevance propagation for explaining deep neural network decisions in MRI-based alzheimer’s disease classification. Frontiers in Aging Neuroscience, 11, jul 2019.

[24] Erico Tjoa, Guo Heng, Lu Yuhao, and Cuntai Guan. Enhancing the extraction of interpretable information for ischemic stroke imaging from deep neural networks. 2019.

[25] Michel van Kempen, Stephanie S. Kim, Charlotte Tumescheit, Milot Mirdita, Johannes Söding, and Martin Steinegger. Foldseek: fast and accurate protein structure search. bioRxiv, 2022.

[26] Sean Whalen, Jacob Schreiber, William S. Noble, and Katherine S. Pollard. Navigating the pitfalls of applying machine learning in genomics. Nature Reviews Genetics, 23(3):169–181, November 2021.

[27] Janani Durairaj, Joana Pereira, Mehmet Akdel, and Torsten Schwede. What is hidden in the darkness? Characterization of AlphaFold structural space. bioRxiv, 2022.

[28] Noelia Ferruz, Florian Michel, Francisco Lobos, Steffen Schmidt, and Birte Höcker. Fuzzle 2.0: Ligand binding in natural protein building blocks. Frontiers in Molecular Biosciences, 8, August 2021.

[29] Craig O. Mackenzie, Jianfu Zhou, and Gevorg Grigoryan. Tertiary alphabet for the observable protein structural universe. Proceedings of the National Academy of Sciences, 113(47), November 2016.

[30] Diederik P Kingma and Max Welling. Auto-Encoding Variational Bayes. arXiv, ec 2013.

